# Neutrophils exhibit distinct migration phenotypes that are regulated by transendothelial migration

**DOI:** 10.1101/2024.10.17.618860

**Authors:** Amy B. Schwartz, Adithan Kandasamy, Juan C. del Álamo, Yi-Ting Yeh

**Affiliations:** Department of Mechanical and Aerospace Engineering, University of California San Diego, La Jolla, CA; Department of Mechanical Engineering, University of Washington, Seattle, WA; Center for Cardiovascular Biology, University of Washington, Seattle, WA; Institute of Stem Cell and Regenerative Medicine Cardiovascular Biology, University of Washington, Seattle, WA

## Abstract

The extravasation of polymorphonuclear neutrophils (PMNs) is a critical component of the innate immune response that involves transendothelial migration (TEM) and interstitial migration. TEM-mediated interactions between PMNs and vascular endothelial cells (VECs) trigger a cascade of biochemical and mechanobiological signals whose effects on interstitial migration are currently unclear. To address this question, we cultured human VECs on a fibronectin-treated transwell insert to model the endothelium and basement membrane, loaded PMN-like differentiated HL60 (dHL-60) cells in the upper chamber of the insert, and collected the PMNs that crossed the membrane-supported monolayer from the lower chamber. The 3D chemotactic migration of the TEM-conditioned PMNs through collagen matrices was then quantified. Data collected from over 50,000 trajectories showed two distinct migratory phenotypes, i.e., a high-persistence phenotype and a low-persistence phenotype. These phenotypes were conserved across treatment conditions, and their existence was confirmed in human primary PMNs. The high-persistence phenotype was characterized by more straight trajectories and faster migration speeds, whereas the low-persistence one exhibited more frequent sharp turns and loitering periods. A key finding of our study is that TEM induced a phenotypic shift in PMNs from high-persistence migration to low-persistence migration. Changes in the relative proportion of high-persistence and low-persistence populations correlated with GRK2 expression levels. Inhibiting GRK2 hindered the TEM-induced shift in migratory phenotype and impaired the phagocytic function of PMNs. Overall, our study suggests that TEM-mediated GRK2 signaling primes PMNs for a migration phenotype better suited for spatial exploration and inflammation resolution. These observations provide novel insight into the biophysical impacts of TEM that priming PMNs is essential to conduct sentinel functions.

## Introduction

Polymorphonuclear neutrophil (PMN) migration to sites of injury or infection is a primary component of innate immunity and inflammatory responses (1, 2). Upon encountering the signals associated with injured or inflamed tissue, circulating PMNs escape blood vessels by adhering to and probing inflamed vascular endothelial cells (VECs), breaching endothelial cell-cell junctions, and transmigrating across the endothelium before infiltrating the surrounding tissue (3). This process, known as trans-endothelial migration (TEM), is a rate-limiting step in the immune response (1). Transmigrated PMNs efficiently navigate 3D tissue-like environments to reach their targets (4–6). The interactions between PMNs and VECs during TEM modulate several PMN characteristics, including their life span and effector function (7, 8). However, while the initial recruitment of PMNs has been extensively characterized (1, 3), the effect of TEM on their subsequent interstitial migration behavior, and the correspondence between resulting migration phenotype and effector function, remain largely unknown.

There are two mechanisms whereby PMNs can traverse the VEC monolayer and exit the circulation: paracellular TEM through the junctions between VECs and transcellular TEM through thin portions of the endothelial cell body (1). The paracellular route is the most common, comprising over 90% of transmigration events (9). Paracellular TEM is facilitated by interactions with many endothelial-expressed surface adhesion molecules, including vascular endothelial cadherin (VE-cadherin), platelet/endothelial-cell adhesion molecule 1 (PECAM1), various junctional adhesion molecules, and other selective adhesion molecules such as ICAM-1 (3, 10). TEM is also known to induce PMN surface expression of the β1-integrin family and β2-integrins, such as lymphocyte function-associated antigen 1 (LFA1) and macrophage-1 antigen (CD11b) (1). Mechanically, PMNs extend invadosome-like protrusions to dynamically probe the underlying endothelium and potentially identify the path of least resistance (11). Subsequently, they actively contract over the VECs, leading to junctional gap openings, and force themselves through the gaps (12). VECs also play an active role in PMN trafficking, forming actin-rich ICAM-1 projections known as “docking structures” that help to anchor and embrace the leukocyte and have been speculated to facilitate extravasation by generating contractile stresses to pull PMNs into and across the VEC monolayer (13, 14). Because PMN sizes can be more than 20 times greater than the size of endothelial cell-cell junctions (∼10 µm vs. ∼0.5 µm), transmigrating through VEC junctions can cause unusually high PMN deformations (15). Mechanical constriction at VEC junctions has been suggested to influence PMN transmigration and subsequent interstitial migration towards infection sites (7).

*In vivo*, the extravascular space is a complex, highly dynamic, and densely packed environment composed of collagen fibers, glycoproteins, and other ECM fibers. As such, transmigrated PMNs need to negotiate crowded spaces with varying pore sizes to reach the site of inflammation (6, 16, 17). In 3D environments, PMNs utilize integrin-independent migration, relying on actin cytoskeletal-network expansion to generate protrusive forces along the leading edge, coupled with myosin-II dependent contractility to bring up the rear (18, 19). To perform efficient pathfinding in 3D, PMNs alternate frequent turns with directional persistence to steer themselves toward the chemotactic gradient (4). This alternance involves intermittent, integrin-mediated periods of traction force exertion (20). Cell migration in 2D is typically modeled as a persistent random walk (PRW), which depends on two critical motility parameters: migration speed and persistence (21–24). However, the complexity of cell-ECM interaction in restrictive 3D environments introduces modeling difficulties (25). Also, single cells moving in 3D can rapidly transition between slow and fast migration states, making it necessary to contemplate time-varying parameters in random walk models (26). In addition to these dilemmas, PMNs can exhibit distinct phenotypic diversity and plasticity in crucial functions, including their migratory behavior (27–30). For example, successful clearance of bacterial infection depends on efficient PMN migration into the infected tissues, including a stop signal that helps resolve inflammation (31). However, phenotypic diversity in migration is rarely considered in experimental analysis and remains poorly understood.

The early response of PMNs to a chemoattractant gradient increases migration speed and persistence towards the chemoattractant focus. In contrast, these migratory parameters are suppressed in late response phases as high chemoattractant concentrations desensitize surface G-protein coupled receptors (GPCRs) on the cell surface (32–34). The multifunctional GPCR kinase GRK2 was recently linked to PMN swarm resolution via receptor desensitization, with GRK2-deficient PMNs also demonstrating impaired bacterial clearance by inefficiently picking up and ingesting microbes from bacteria clustering (31). In addition to its classical role in modulating GPCR signaling through receptor internalization and recycling, GRK2 is known to function as a signaling hub via phosphorylation and scaffolding interactions with myriad non-GPCR partners (35). Of note, GRK2 has been shown to negatively regulate immune response via direct association with the MAPK kinase MEK (36) and to phosphorylate the membrane-cytoskeletal linkers ezrin (37) and radixin (38), suggesting a direct link between GRK2 expression and actin cytoskeleton reorganization. Based on these results, GRK2 presents an intriguing link between PMN migration phenotype and effector functionality.

This work’s central hypothesis is that the biomechanical PMN-VEC interactions associated with TEM modulate PMN functions to promote the efficient identification and elimination of pathogens. To address this hypothesis, we employed an experimental assay with two subsequent stages. The first stage is a transwell-based TEM model, and the second stage is a physiologically relevant 3D collagen directional migration assay. Using this in vitro model, we characterized the migratory and functional changes induced by TEM of nearly 50,000 individual dHL-60 and primary PMN cells. We used sequential and variational Bayesian inference algorithms to identify two naturally occurring, phenotypically distinct migratory subpopulations conserved across treatment conditions. The first is a high-persistence phenotype characterized by a fast, directed migration toward chemoattractant sources. The second is a low-persistence phenotype characterized by a slower, more random migration. The relative prevalence of these phenotypes is modulated by TEM-mediated changes in PMN expression of GRK2. These findings show a critical role for TEM in determining the subsequent PMN interstitial migration phenotypes, demonstrating that VECs can modulate PMN migration to promote immune function.

## Results

### TEM mtysodulates transcription and protein expression of chemokine receptors and increases phagocytic function

To model the impact of PMN trans endothelial migration (TEM) on subsequent interstitial 3D migration, we developed a two-stage *in vitro* assay that combines a commercial transwell migration assay and a previously described 3D-migration chamber (4) (Figures 1A and 2A). We performed real-time quantitative PCR (RT-qPCR; Figure 1B) and flow cytometry (FCS; Supplementary Figure 1A) to compare intracellular gene expression and cell surface protein expression, respectively, of selected chemokine receptors, surface integrins, and glycoproteins before and after cells crossing the VEC-coated transwell insert (**ETW**-conditioning). These data collectively demonstrate upregulation of CD11b, ICAM-1, and FPR1 (for RT-qPCR, ICAM-1: p = 0.0024; FPR1: p = 0.0172) in conjunction with downregulation in CXCR2 (for RT-qPCR, p = 0.003) and decreased cell surface expression of CXCR1 (for RT-qPCR, p > 0.9999). These changes are all qualitatively consistent with PMNs that have undergone TEM *in vivo* (39), suggesting that TW crossing altered the expression profile of dHL-60 cells in a similar manner.

**Figure 1.**
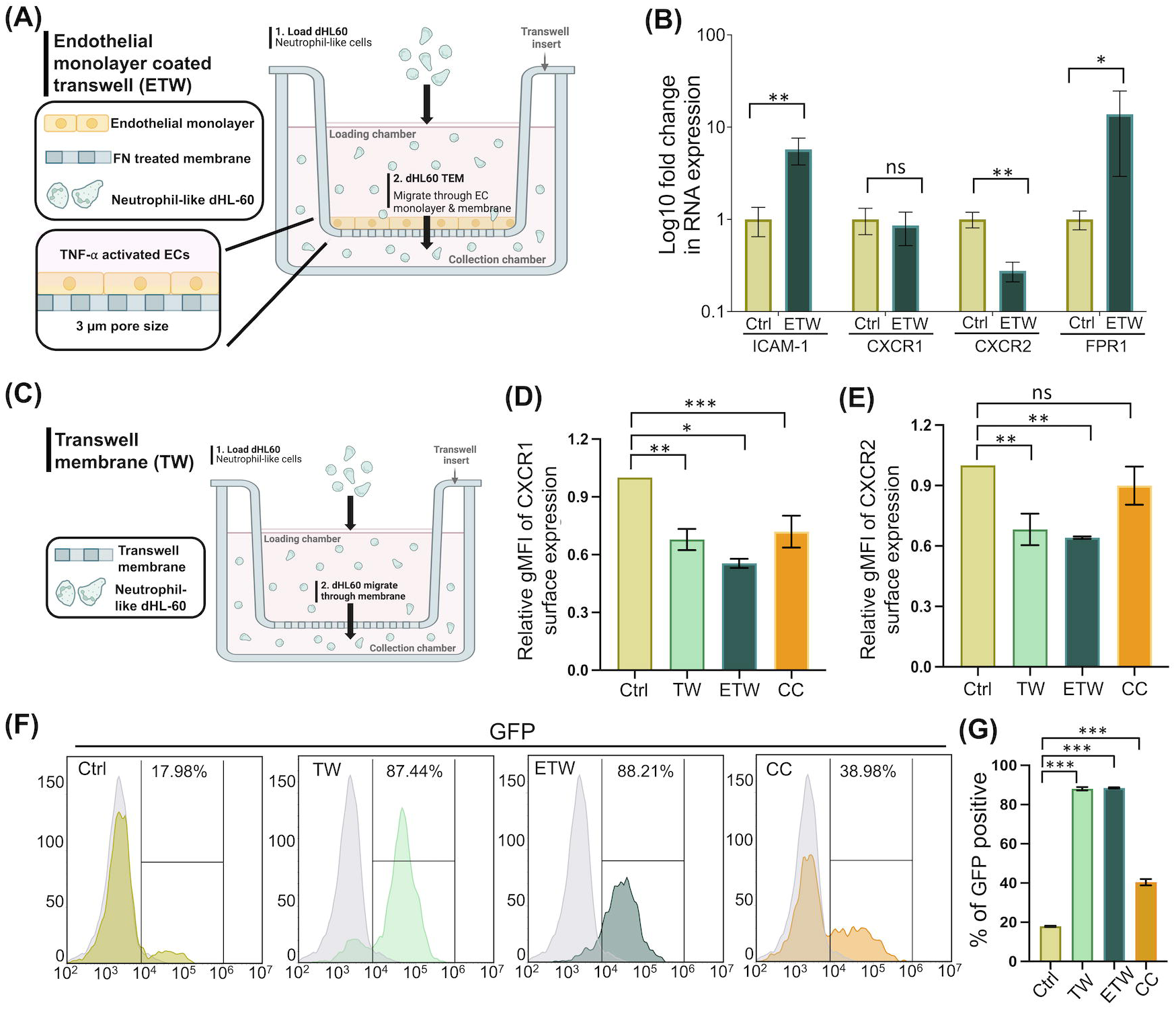
TEM process alters neutrophil phenotypes associating with the loss of directional motility and better phagocytosis functions. (A) Schematic of the Transwell-based treatment assay employed to replicate the physiological effects of TEM on PMN physiology. VECs are seeded on the polycarbonate membrane of a TW insert and grown to confluence and activated with TNF-α before day 5 DMSO-differentiated HL-60 cells are loaded into the upper chamber. PMNs that transmigrate through the VEC monolayer and TW membrane are collected from the lower chamber for use in further experiments. (B) RT-qPCR results of gene expressions in dHL-60 cells subjected to static control or ETW-crossing cells. Values are expressed in terms of log fold change compared to control. Error bars were calculated for ΔC_T_ values. n=3, \**p* < 0.05, \*\**p* < 0.01 Krystal Wallis test. (C) TW condition where dHL-60 cells transmigration through the Transwell membrane with pore diameter of 3 µm. (D-E) FCS results of cell surface CXCR1 and CXCR2 expressions in control, TW, ETW and CC-conditioned dHL-60 cells. Fluorescent expression levels of CXCR1 and CXCR2 on TW, ETW and CC-conditioning cells are relative to levels of the control dHL-60 cells. (F) Flow cytometry demonstrating the signal intensity of GFP in dHL-60 cells. The gray color intensity is from dHL-60 cells without interacting with zymosan, showing the basal level. (G) Quantitative analysis of GFP positive cells after 3 hours incubation with pHrodo zymosan. (D-F) *p* < 0.05, \*\**p* < 0.01, \*\*\**p* < 0.001 (versus control group) using a one-way ANOVA with Dunnett’s post multiple comparison test. n = 3.

**Figure 2.**
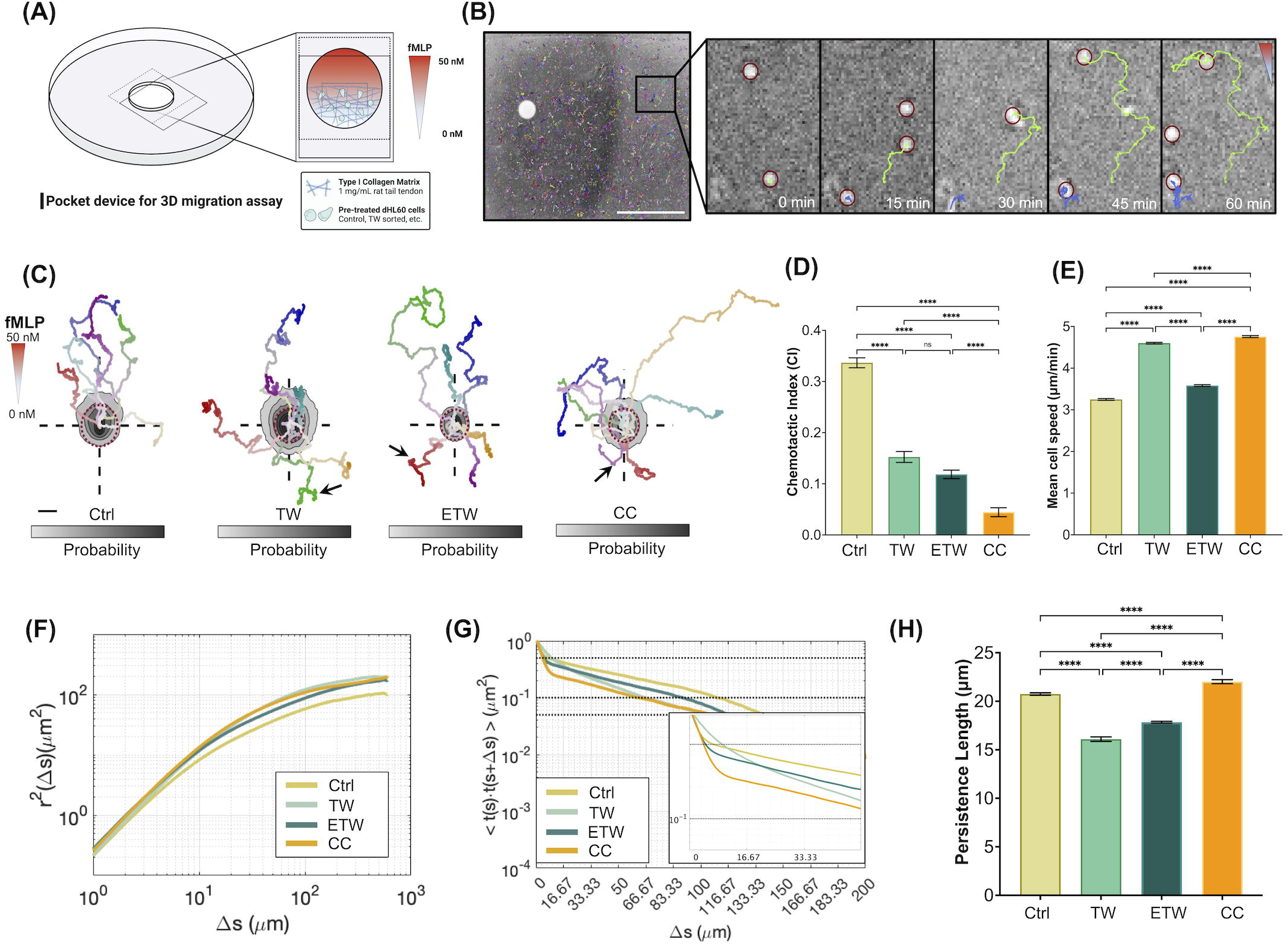
TEM induces heterogenous neutrophil motility in 3D accounting for overall reduced area exploration and chemotactic migration. (A) Schematic of pocket device for *in vitro* 3D directed neutrophil migration assays. (B) Sample image of cells migrating in a pocket device with Trackmate cell tracking results overlayed. Inset shows the time sequence for a chemotaxing cell, with tracked trajectory overlayed in green. Scale bar, 1000 µm. (C) Probability density maps of aggregated trajectory endpoints are shown for dHL-60 cells in each of the four conditions with representative sample tracks overlayed. Sample track color saturation increases with time. The dashed red reference circle superimposed on each inset figure represents unbiased random motion. fMLP concentration increases linearly in the positive y direction. Black arrows show the large angle turns. Scale bar, 20 µm. (D) Population average chemotactic index (CI). (E) Population average mean cell speeds in four treatment conditions in the presence of fMLP gradient. Error bars represent 95% confidence interval. All pairwise conditions showed statistically significant differences besides those labeled n.s. (not significant) in (D-E). \*\*\*\**P* < 0.0001 according to the one-way ANOVA Tukey’s test. (F) Mean squared displacements (MSDs) versus distance separation along cell trajectories, *r*^2^(Δ*s*). (G) Tangent vector autocorrelations versus distance traveled, (*Ctt* ((Δ*s*)), for *f*MLP condition, of four treatment conditions. The inset represents the enlarged portion of the initial decay. (H) Migratory persistence length, L_P_. Values reflect the characteristic length of the slow decaying component of the double exponential fit, with error bars calculated via bootstrapping (see Methods). \*\*\*\**P* < 0.0001 according to the Brown-Forsythe ANOVA test.

PMNs utilize their G-protein-coupled chemokine receptors (GPCRs) to detect chemoattractants (40). Two receptors, CXCR1 and CXCR2, have been reported to coordinate PMN directional motility through differential surface expression levels, promoting PMN clustering at the infection site and facilitating the posterior self-resolution of PMN clusters (34). The down-regulation of CXCR1 and CXCR2 gene and surface expressions with ETW crossing prompted us to investigate how TEM impacted PMN directional migration and phagocytic function. Thus, we conducted additional experiments using dHL-60 cells co-cultured with VECs without transmigration (co-culture condition, **CC**) and loaded into unmodified transwell inserts (transwell only condition, **TW**) to decouple the contribution of VECs and the cellular deformation experienced by PMNs squeezing through narrow transwell membrane pores (Figure 1C). We performed flow cytometry to detect surface expressions of CXCR1 and CXCR2 in these four conditions, finding that the surface expressions of both CXCR1 and CXCR2 decrease significantly in TW- and ETW-conditioned cells (Figures 1D-E). In contrast, the expression of CXCR2 did not decrease significantly after CC conditioning. Taken together, these results suggest that the surface expressions of PMN chemokine receptors are mechanosensitive and that biochemical interactions with VECs may regulate surface trafficking of CXCR1 and CXCR2 through different mechanisms.

It is important to note that the CXCR2 fluorescence intensity profiles exhibited a marked shift to low values for TW- and ETW-conditioned cells (Supplementary Figure 1B). The large magnitude of this shift allowed us to discard the possibility that the changes in CXCR2 with respect to controls were the result of a selection process, i.e., that cells with low CXCR2 expression values were more successful at crossing the transwell membrane pores, because cells with such low levels of CXCR2 expression did not originally exist in the control and CC-treated groups. Therefore, we concluded that these changes were a phenotypic change induced by TW- and ETW-conditioning.

Next, we used pHrodo GFP-conjugate zymosan to assess changes in PMN phagocytosis ability following these treatments. Zymosan particles do not fluoresce outside of the cells or inside of the cell cytosol due to these environments’ neutral pH. However, when phagocytes internalize the particles into the acidic environments of endosomes and lysosomes, the GFP dye fluoresces brightly, reporting bacterial ingestion into PMN phagosomes. Flow cytometry results showed almost 90% of TW- and ETW-conditioned cells were GFP-positive for bacterial ingestion (Figure 1F). In comparison, only 40% of CC-conditioned cells and 20% control cells were GFP-positive (Figure 1G). These data demonstrate that mechanical stimuli in the TEM process can activate the PMN phagocytic machinery.

### The persistence of PMN migration through 3D collagen matrices is heterogeneous

To study the impact of TEM on the subsequent PMN migration through 3D matrices, we tracked and analyzed the motion of approximately 50,000 control, CC, TW-, and ETW-conditioned cells in 1 mg/mL collagen under an fMLP chemoattractant gradient as previously described (4) (Figures 2A-B). Figure 2C shows representative trajectories for chemotaxing dHL-60 cells in each condition superimposed onto probability density maps of their endpoint positions. A red dashed circle representing non-chemotactic random migration is included for reference. Control cells migrated directionally up the fMLP gradient as expected (4), but cells from all other conditions described omnidirectional trajectories. In particular, the cells from the TW, ETW, and CC appeared to have an increased propensity to migrate against the gradient and to incorporate large-angle turns in their migration (see arrows in Figure 2C), similar to those reported in late phases of the immune response when PMNs are desensitized to the chemoattractant (31). As a result of these behaviors, TW, ETW, and CC cells had significantly lower mean chemotactic indices than controls (Figure 2D) even if their mean speed was higher (Figure 2E). These results are consistent with the changes in CD11b, CXCR1, and CXCR2 reported in the previous section, further suggesting that TEM predisposes PMNs to adopt a more active, exploratory interstitial migration phenotype.

To quantify in more detail how TW, ETW, and CC conditions impacted the cell trajectories, we computed the mean squared displacements (*r*^2^or MSD, Figure 2F) and autocorrelation function of the vector tangent (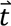) to the cell trajectory (*C_tt_*, Figure 2G) versus distance traveled (Δ*s*). Because the MSD plot is obtained by integrating the *C_tt_* plot twice with respect to Δ*s*, it experiences quadratic growth (*r*^2^∼ Δ*s*^2^) near the origin. As *C_tt_* falls when Δ*s* increases, the MSD plot tapers to behave linearly (*r*^2^ ∼ Δ*s*) for very long distances. This attenuation in slope reflects the loss of overall migratory persistence when migrating in 3D across large distances, as cells encounter obstacles in the microenvironment and need to circumvent them. Despite their slight differences, the MSD of TW, ETW, CC, and control cells were similar (Figure 2F), suggesting all cell trajectories had similar persistence at the population level, even if control cells were persistent and also aligned with the chemoattractant gradient while TW, ETW, CC cells were persistent and omnidirectional.

Inspection of *C_tt_* revealed more complex behavior with two distinct exponential decays, evidenced as two straight segments in each curve of the linear-logarithmic plot in Figure 2G. In all conditions, there was steep initial decay in *C_tt_* over displacements of 0 µm to 10 µm followed by a shallower decay over much longer distances. The existence of two regimes in *C_tt_* suggests that the studied cell populations may exhibit heterogeneous migratory behavior in terms of directional persistence, with the steeper segments corresponding to a less motile phenotype than the shallow segments. This possibility is evaluated in detail in the next section. We estimated the persistence length (*L_p_*) of cell trajectories in each condition by performing an exponential fit to the shallower decay segment (see Materials and Methods). The results of this analysis (Figure 2H) indicate that TW- and ETW-conditioned cells experienced a modest albeit significant decrease in *L_p_*, consistent with a more exploratory migratory behavior.

### PMNs exhibit distinct migratory phenotypes whose prevalence is modulated by TEM

Motivated by the results in the previous section, we investigated the existence of distinct migratory phenotypes in PMN cell populations and how TEM affects their relative frequencies. We fitted each cell’s trajectory to a random walk model with time-dependent parameters *q*_*t*_ and *a*_*t*_ (see Materials and Methods) (26). In this model, *q*_*t*_ represents the persistence of the motion and *a*_*t*_ its activity, or instantaneous speed of random motion. We then plotted probability density contours of (*q*_*t*_, *a*_*t*_) for each of the four conditions studied: TW, ETW, CC, and control (Figures 3A-D). The probability contours obtained for control cells were clearly bimodal (Figure 3A), revealing two distinct migratory phenotypes. The most abundant one was formed by cells of high persistence and activity, whereas the other consisted of significantly less persistent and slightly less active cells. We named them high-persistence and low-persistence phenotypes and observed their presence in the other conditions. In TW-conditioned cells, the high-persistence phenotype was very dominant, and the low-persistence one was virtually nonexistent (Figure 3B). Conversely, the high-persistence phenotype was less abundant, and the low-persistence phenotype was more prevalent in ETW-conditioned and CC-conditioned cells than in controls (Figure 3C and 3D, respectively).

**Figure 3.**
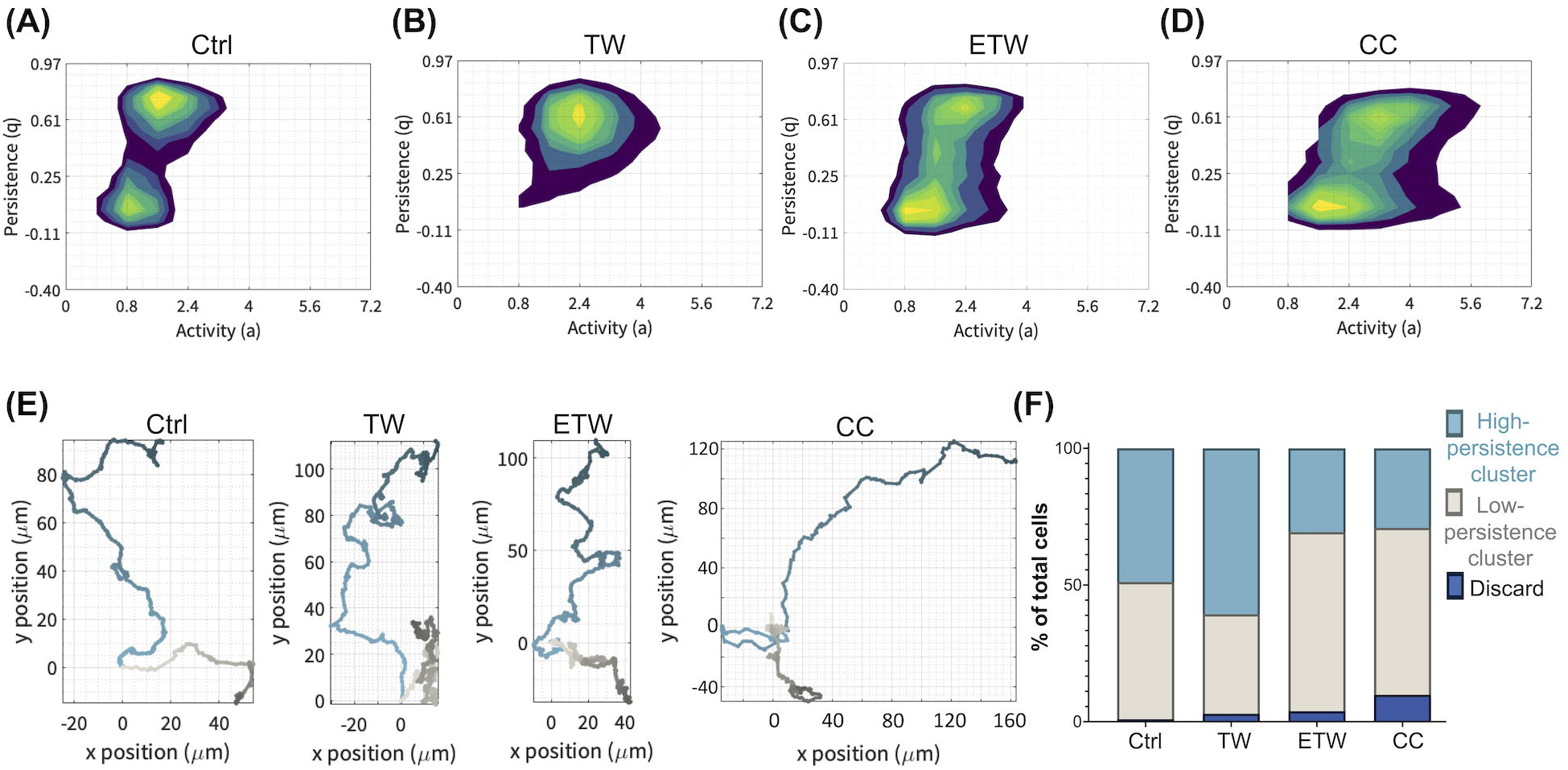
Mapping individual cell trajectories to time-dependent activity and persistence parameters of a heterogeneous persistence random walk reveals distinct migratory subpopulations. (A-D) Probability density contour plots for motility modes, comprised of paired activity and persistence values, for each of the four principal treatment conditions. Contours are plotted for 5, 10, 20, 40, 60, 80, and 90% of total population density for each condition. (E) Representative sample tracks taken from the high-(blue) and low-(grey) persistence clusters for each of the four conditions. Color darkens along the course of each trajectory to indicate time progression. fMLP gradient concentrations increases in direction of increasing y. (F) Relative prevalence of each migratory subpopulation for each of the four conditions, expressed as percentage of total population for each condition. Low-persistence cluster includes non-motile cells.

To confirm the existence of distinct motility phenotypes, we ran a quasi-unsupervised clustering algorithm rooted in variational Bayesian inference for a Gaussian Mixture Model (VB-GMM, see Materials and Methods). The VB-GMM algorithm was tasked with identifying four clusters in (*q*_*t*_, *a*_*t*_) space from the pooled distribution of all the studied dHL-60 cells (Supplementary Figure 2A). Visual inspection of the cell trajectories belonging to the least abundant cluster (extremely high activity levels, <10% of the total cell population) revealed that these trajectories had obvious cell tracking errors. As such, this subgroup was labeled “Discard”, and its tracks were excluded from further analysis. The remaining two clusters identified by the VB-GMM algorithm were neatly aligned with the two modal peaks in the probability contours of Figures 3A-D and were, therefore, similarly denoted as high- and low-persistence clusters (Supplementary Figure 2B).

Figure 3E shows examples of trajectories belonging to each cluster for each of the conditions studied, confirming that high-persistence trajectories were long and relatively straight, whereas low-persistence trajectories described numerous turns and covered less distance. Figure 3F quantifies the relative prevalence of each motility cluster across all studied conditions, indicating that control cells had nearly an even split between low- and high-persistence trajectories (48.9% high, 50.2% low) and that this split changed for TW-conditioned, ETW-conditioned, and CC-conditioned cells. Consistent with the probability maps described above, the high-persistence trajectories became more frequent in TW-conditioned cells (60.7% high, 36.4% low). Likewise, the low-persistence trajectories became more frequent in ETW-conditioned cells (30.6% high, 65.5% low) and CC-conditioned cells (29.1% high, 61.0% low).

These experiments demonstrate that dHL-60 cells exhibit a heterogeneous migratory behavior characterized by two distinct phenotypes: a direct migration phenotype and a more wandering phenotype. Our data also suggests that TEM predisposes dHL-60 cells to switch from the direct phenotype to the wandering phenotype. Taken together, our results support that signaling induced by interactions with the VECs plays a dominant role in promoting the interstitial migratory phenotype of dHL-60 cells after TEM. In our experiments, VEC-dHL60 contact eclipsed the phenotypic change induced by mechanical deformation alone (TW condition).

### Primary human PMNs display high- and low-persistence migratory phenotypes as observed in dHL-60 cells

DMSO differentiated HL-60 cells are often employed as a PMN model as their migration and function are qualitatively similar to primary PMNs (41). To elucidate if the existence of two migratory phenotypes represents primary PMN behavior, we repeated the 3D interstitial migration experiments and subsequent analyses with primary PMNs derived from whole human blood (Materials and Methods). The probability density contours of (*q*_*t*_, *a*_*t*_) for primary PMNs, shown in Figure 4A, agreed qualitatively with those obtained from dHL-60 cells, overlaid for reference in the figure. More specifically, PMNs exhibited a low-persistence mode that co-localized with the low-persistence mode of dHL-60 cells in the (*q*_*t*_, *a*_*t*_) space. Besides, the primary PMN probability density map displayed a long horizontal plateau with similar persistence values and equal or higher activity values than those of the dHL-60 high-persistence mode. We identified this plateau as the primary PMN high-persistence mode.

**Figure 4.**
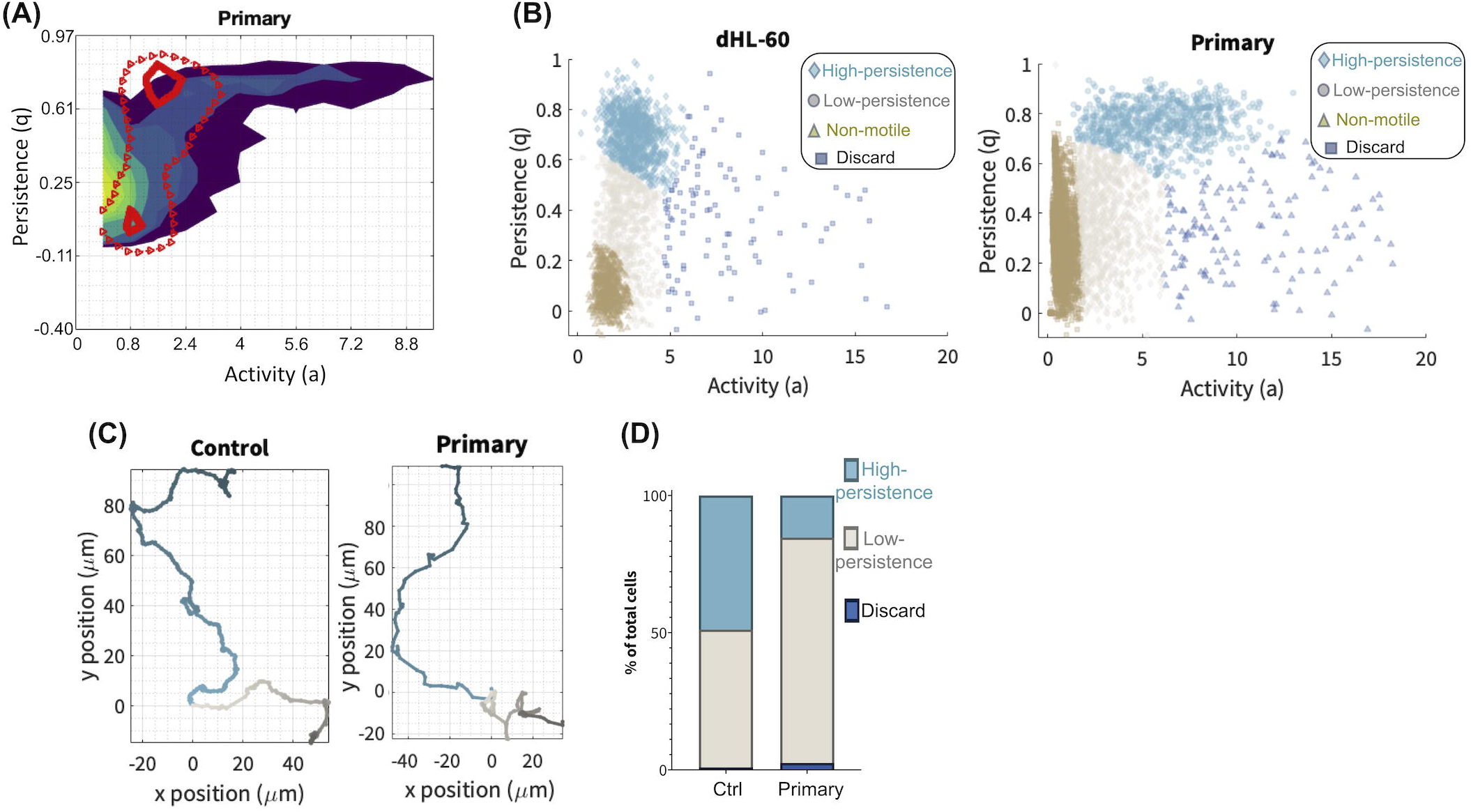
Distinct motility subpopulations are conserved in PMNs. (A) Probability density contour plot for paired activity/persistence motility modes of PMNs, with contours plotted for 5, 10, 20, 40, 60, 80, and 90% of total population density. 30% and 90% contours of control dHL-60 cells (red triangles) are overlayed for reference. (B) Cluster map with cell averaged 3D persistent random walk parameters (a,q) in PMNs and dHL-60 cells. Variational Bayesian inference for a Gaussian Mixture Model (VB-GMM) was performed (see Methods). The motility clusters comprising both the high-persistence (↑q) and low-persistence (↓q) phenotypes. Based on the sample tracks, the non-motile (brown) is distinguished from a low motility ↓q (gray) cluster for further analysis of proportions by VB-GMM. (C) Representative sample tracks taken from the high-(blue) and low-(grey) persistence clusters for control dHL-60 cells and PMNs. Color darkens along the course of each trajectory to indicate time progression. fMLP gradient concentrations increases in direction of increasing y. (D) Relative prevalence of each migratory subpopulation for PMNs and control dHL-60, expressed as percentage of total population for each condition. Low-persistence cluster includes non-motile cells.

When applied to the pooled dataset of tracked primary PMN trajectories, the VB-GMM algorithm independently identified distinct high-persistence and low-persistence clusters (Figure 4B). Moreover, these clusters occupied similar regions of the (*q*_*t*_, *a*_*t*_) space as dHL-60 cells, even if primary PMNs were skewed slightly toward higher activity values overall. Example trajectories obtained from the low- and high-persistence clusters had similar appearances in dHL-60 cells and primary PMNs (Figure 4C-D). On the other hand, the relative frequency of high-persistence migration was lower in primary PMNs s (15.4%) than in dHL-60 cells (48.9%), as shown in Figure 4E.

Summary migration statistics were computed separately for the high- and low-persistence clusters of primary PMNs to permit a quantitative fine-grained comparison with dHL-60 cells. For quantifying effective migration statistics, we excluded the non-motile cells (Supplementary Figure 2A). The mean cell speeds of both cell types in the high-persistence cluster were significantly different, with primary PMNs in the control condition moving approximately twice faster than dHL-60 cells in all conditions (Figure 5A). In the low-persistence cluster, the cell speeds of control condition of dHL-60 cells and primary PMNs still showed statistically significant differences, but the differences in magnitude were modest (Figure 5B). Likewise, TW, ETW, and CC conditioning increased the speed of dHL-60 cells in both clusters, but these differences were considerably smaller than the differences between clusters. When analyzing the chemotactic efficiency of cells belonging to each subgroup (Figure 5C-D), we found that TW, ETW, and CC conditioning caused a >50% reduction in CI in both the high- and low-persistence clusters, bringing the net CI to near zero values (<0.1) in the low-persistence cluster. Of note, ETW and CC conditioning brought the chemotactic efficiency of dHL-60 cells to values similar to primary PMNs. These data suggest that VEC interactions are crucial for dHL-60 cells to better recapitulate primary cell characteristics.

**Figure 5.**
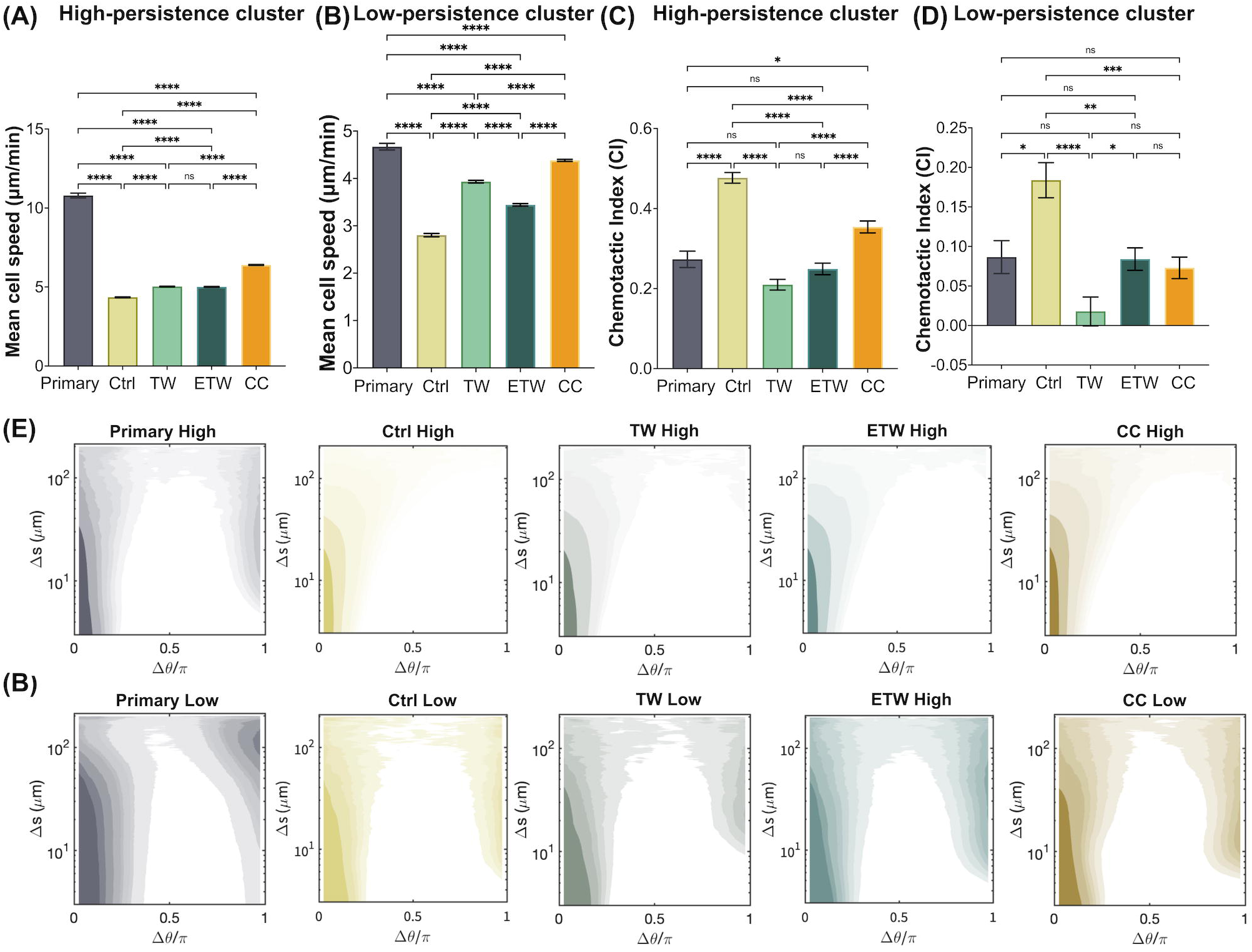
Quantitatively characterizing migration statistics reveals the dHL-60 low persistence subpopulation replicates key features of low persistence PMN behavior. Descriptive statistics calculated for PMNs as compared to dHL-60 cell with treatment conditions for cells assigned to high- and low-persistence subpopulations: (A-B) instantaneous mean cell speed, (C-D) chemotactic index. (A-D) \**p* < 0.05, \*\**p* < 0.01, \*\*\**p* < 0.001, \*\*\*\**P* < 0.0001 according to the one-way ANOVA Tukey’s test. (E-F) Probability distribution maps of turning angles (Δ*θ*) conditional to separation distance (Δ*s*), *p*(Δ*θ*|Δ*s*) of the high-persistence cluster (E) and the low-persistence cluster (F) in PMNs and dHL-60 cells of the four conditions.

### TEM-induced low-persistence migration involves zigzagging

Previous studies (4, 25) have uncovered hidden features of 3D cell migration using the probability distribution of turning angles conditional to the distance traveled along each cell’s trajectory, *p*(Δ*θ*|Δ*s*). Thus, we computed this distribution for the high- and low-persistence trajectories of primary PMNs and dHL-60 cells in all conditions studied. The *p*(Δ*θ*|Δ*s*) maps from the control dHL-60 cells belonging to the high-persistence cluster (Figure 5E) displayed a narrow peak around *Δ*θ* = 0* for short distances Δ*s*, which broadened as Δ*s* increased. This behavior has been attributed to cells incorporating turns in their trajectories to circumvent environmental obstacles, leading to a progressive loss of directional memory with increasing spatial separation (4). Highly motile dHL-60 cells in the TW, ETW, and CC conditions exhibited similar *p*(Δ*θ*|Δ*s*) distributions (Figure 5E).

Cells assigned to the low-persistence cluster (Figure 5F) exhibited markedly different distributions of turning angles. The low-Δ*s* peak around *Δ*θ* = 0* was broader than in the high-persistence cluster, suggesting a propensity to describe slightly less straight segments at short distances compared to cells in the high-persistence cluster. Moreover, a second prominent peak appeared around *Δ*θ* = p* for Δ*s* > 10 mm, indicating an increase in the frequency of ≈180-degree turns. Inspection of individual migration tracks suggest that this occurs because cells began implementing abrupt changes of direction alternating with relatively straight segments, i.e., a zigzagging behavior (42) (Supplement Figure 3). The turning angle distributions for cells in the TW, ETW, and CC conditions had similar bimodal features (Figure 5F).

The similarities in the *p*(Δ*θ*|Δ*s*) of control, TW-, ETW-, and CC-conditioned cells within each cluster confirm that these motility clusters represent distinct and well-defined migration phenotypes that are conserved across dHL-60 treatment conditions. Importantly, the bimodal turning angle distribution seen in dHL-60 cells was also observed in primary PMNs and was more prominent in the low-persistence cluster (Figure 5E-F). Zigzagging migration patterns have been observed in PMNs *in vivo* and have been proposed to help find sparsely distributed targets (42). Our data suggests these phenotypes, particularly the low-persistence phenotype, are preserved across primary PMNs and dHL-60 cells with high level of detail in terms of trajectory morphology.

### TEM-mediated leukocyte expression of GRK2 controls PMN’s ability of directional migration and phagocytosis

The existence of distinct migratory phenotypes in PMN migration through 3D matrices, and the modulation of their relative prevalence on TEM-associated external stimuli motivated our investigation of molecular signals involved in this modulation. Recent *in vitro* experiments of murine PMN migration demonstrated that GRK2-deficient PMNs exhibit faster, more persistent migration in response to but irrespective of the direction of externally imposed gradients of the intermediate chemoattractant LTB4 (31). Due to the stark shift in migration phenotype seen in knockdown cells, GRK2 immediately became a candidate molecule to explain the differences in track tortuosity between the high- and low-persistence clusters observed in our experiments.

Figure 6A shows RT-qPCR results for GRK2 expression in the four conditions studied: control, TW, ETW, and CC. Even if these data demonstrate potential for kinase expression rather than actual kinase activity, the statistically significant increases seen in TW- and ETW-conditioned dHL-60 cells are consistent with the observed population-level losses in chemotactic index. Notably, the expression levels in ETW- and CC-conditioned cells were indistinguishable (p > 0.9999), suggesting endothelial interaction is sufficient to induce moderate PMN GRK2 increases. Compared to the stark GRK increase in the TW conditioned, such moderate GRK expressions in ETW-conditioned cells suggest that endothelial interactions may blunt the effect of TEM-mediated mechanical deformation on PMN chemotactic migration.

**Figure 6.**
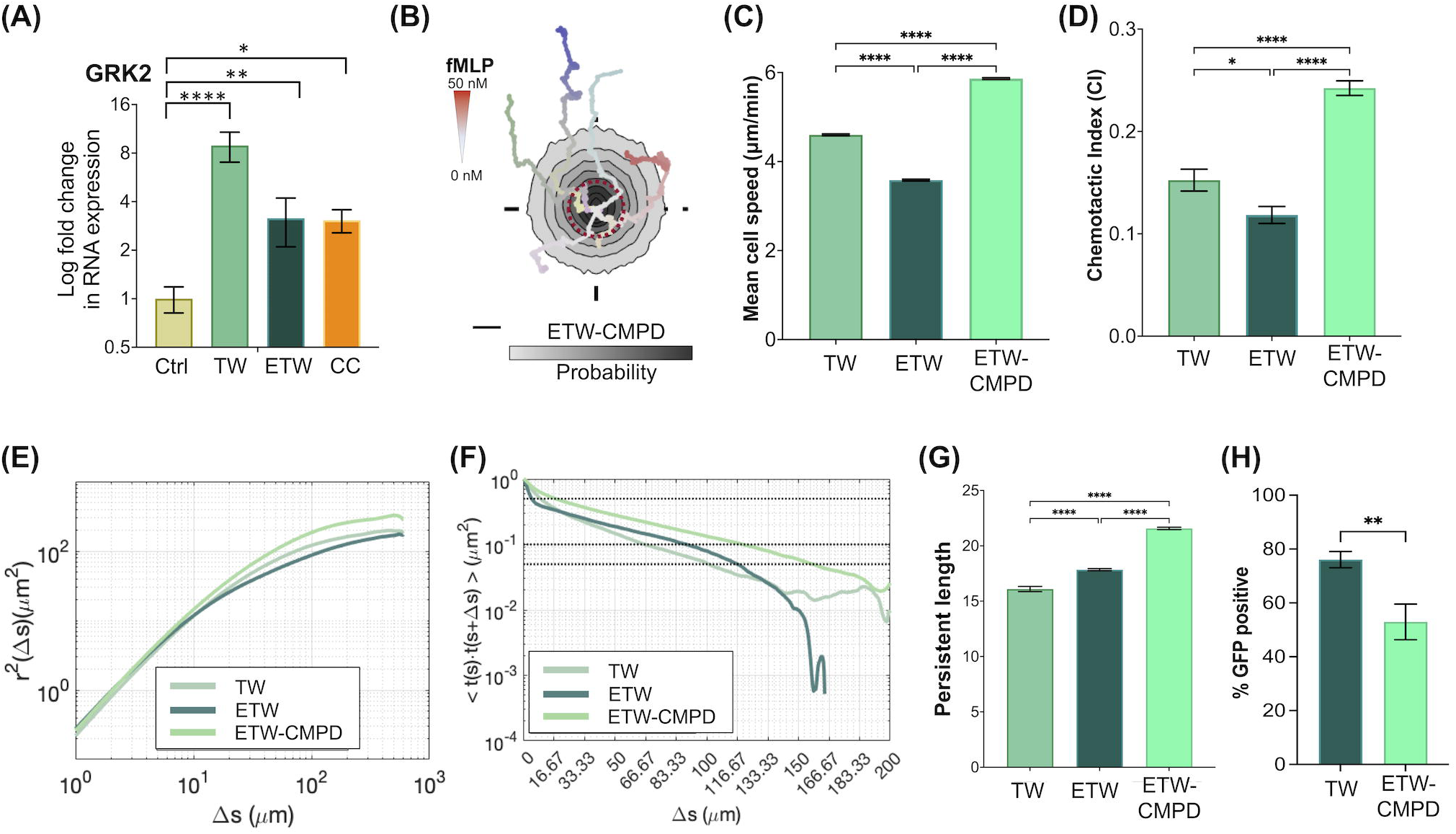
TEM-mediated leukocyte expression of GRK2 controls PMN’s ability of directional migration and phagocytosis. (A) RT-qPCR results indicating changes in internal gene expression potential for GRK2 in all four conditions. Values are expressed in terms of log fold change compared to control. Error bars were calculated for ΔCT values. n = 3. \**p* < 0.05, \*\**p* < 0.01, \*\*\**p* < 0.001 to the one-way ANOVA Dunn’s post t-test. (B) Probability density maps of aggregated trajectory endpoints for ETW + CMPD treated dHL-60 cells with sample tracks overlayed. Sample track color saturation increases with time. The dashed red reference circle superimposed on each inset figure represents unbiased random motion. fMLP concentration increases linearly in the positive y direction. Scale bar, 20 µm. (C) Population level instantaneous mean cell speed, (D) chemotactic index, (E) MSDs and (F) autocorrelation function of the vector tangent to the cell track versus distance traveled (*Ctt* ((Δ*s*)), and (G) migratory persistence length calculated for TW, ETW, and ETW + CMPD treated dHL-60 cells. (H) Flow cytometry demonstrating the signal intensity of GFP in dHL-60 cells. Quantitative analysis of GFP positive cells after 3 hours incubation with pHrodo zymosan. \*\*\**p* < 0.001 (versus control group). n = 3 biological repeats. (C-D) \**p* < 0.05, \*\*\*\**P* < 0.0001 according to the one-way ANOVA Tukey’s test. (G) \*\*\*\**P* < 0.0001 according to the Brown-Forsythe ANOVA test. (H)\*\**P* < 0.01 according to Student’s *t-*test.

We inhibited GRK2 expression in control and ETW-conditioned cells with the potent, selective GRK2 inhibitor CMPD101 and assessed the impact on 3D leukocyte migration before and after ETW-conditioning. CMPD treatment rendered control dHL-60 cells effectively non-motile (Supplementary Figure 4), probably because lower constitutive expression of GRK2 made them unable to perform GPCR trafficking after inhibition. However, and interestingly, the treatment had its anticipated effect on the migration ETW-conditioned cells. Cell trajectories became highly directional and covered more space (Figure 6B), and their mean speed (Figure 6C) and chemotactic index (Figure 6D) increased significantly. Their MSD and *C*_*tt*_ (Figure 6E-F) were suggestive of increased directional persistence, which was confirmed by computing the persistence length *L_p_* and comparing it to that obtained in CMPD-untreated TW- and ETW-conditioned cells (Figure 6G).

In addition to analyzing the migratory behavior of dHL-60 cells after pharmacological GRK2 inhibition, we studied how this inhibition affected the cells’ phagocytic activity, since this function is crucial to the innate immune response. In particular, we evaluated *in vitro* phagocytosis of ETW-conditioned dHL-60 cells with and without CMPD treatment by exposing these cells to zymosan bioparticles (see Materials and Methods). The results from this experiment (Figure 6H) showed that the inhibition of GRK2 activity in the ETW-conditioned cells significantly attenuated TEM-mediated improved zymosan bioparticle ingestion.

### GRK2 controls the TEM-mediated shift from high-persistence to low-persistence interstitial migration

To investigate how GRK2 modulates TEM-mediated changes in migratory phenotype, we tracked and analyzed the trajectories of CMPD-treated, ETW-conditioned dHL-60 cells migrating in our 3D collagen chemotactic device. The resulting probability maps of (*q_*t*_*, *a_*t*_*) were plotted in Figure 7A, together with reference contours from CMPD-untreated, TW-conditioned (grey circles) and ETW-conditioned (red triangle) cells. These data show that CMPD treatment shifted the migratory phenotypic distribution of ETW-conditioned cells towards the distribution observed for CMPD-untreated, TW-conditioned cells. The shift was confirmed by the relative prevalences of low- and high-persistence clusters obtained with the VB-GMM classifier (Figure 7B). Specifically, GRK2 inhibition eliminated nearly 75% of the low-persistence population in ETW-conditioned cells (ETW: 65.50%; ETW-CMPD: 17.45%). When compared with human primary PMNs, the CMPD-treated ETW-conditioned dHL-60 cells had opposite proportions of high- and low-persistence clusters (Figure 7B).

**Figure 7.**
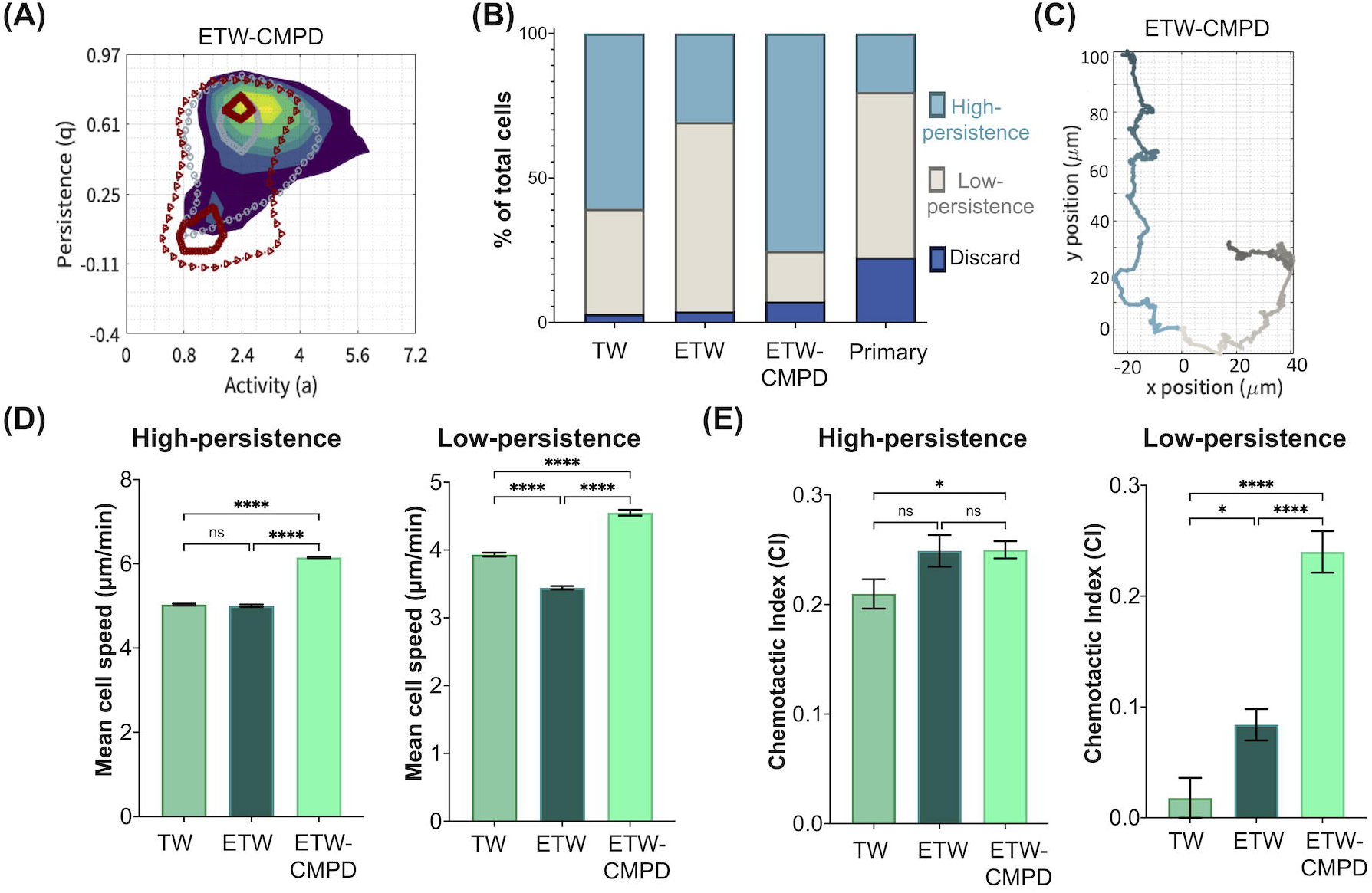
GRK2 controls the TEM-mediated prevalence of low persistence subpopulation. (A) Probability density contour plot for paired activity/persistence motility modes of ETW + CMPD treated cells, with contours plotted for 5, 10, 20, 40, 60, 80, and 90% of total population density. 30% and 90% contours of TW (grey circles) and ETW conditions (red triangles) are overlayed for reference. (B) Relative prevalence of each migratory subpopulation for TW, ETW conditioned, ETW + CMPD, and Primary PMNs, expressed as percentage of total population for each condition. Low-persistence cluster includes the “non-motile” cluster for the comparison. (C) Representative sample tracks taken from the high-(blue) and low-(grey) persistence clusters for ETW + CMPD treated cells. Color darkens along the course of each trajectory to indicate time progression. fMLP gradient concentrations increases in direction of increasing y. (D-E) Descriptive statistics calculated for cells assigned to high- and low-persistence subpopulations of TW, ETW, and ETW + CMPD treated conditions: (D) instantaneous mean cell speed, and (E) chemotactic index. (D-E) \**p* < 0.05, \*\*\*\**p* < 0.0001 using a one-way ANOVA with Dunnett’s test.

Trajectories representative of the high- and low-persistence clusters in CMPD-treated ETW-conditioned cells (Figure 7C) were not strikingly different from those of untreated cells, albeit the low-persistence trajectories in the CMPD-treated cells seemed to zigzag less often. CMPD treatment caused a statistically significant albeit modest mean speed increase in both clusters (Figure 7D). The effect of treatment on the chemotactic index (Figure 7E) was more interesting. While this parameter was not affected by treatment in the high-persistence cluster, it experienced a dramatic increase in the low-persistence cluster. To interpret this result, it is important to keep in mind that the low-persistence cluster in ETW-sorted CMPD-treated cells had low prevalence, so this cluster’s high mean CI might be influenced by having sampled cells from the tail of the more abundant high-persistence cluster. This would be consistent with the high- and low-persistence clusters having similar values of CI after CMPD treatment (Figure 7E). In any case, the main result that holds independent of these methodological details is that cells with low chemotactic efficiency were hardly observed after CMPD treatment.

Altogether, these data show that TEM enhances PMN host-defense function through upregulation of GRK2. The data supports the hypothesis that GRK2 expression is responsible for the tortuous, a chemotactic migration phenotype observed in low persistent cells. Furthermore, CMPD treatment of ETW-conditioned cells recapitulated the migratory mode distribution of TW cells, supporting a role for VEC interaction in modulating GRK2 expression. When considered together, these results implicate GRK2 expression as the controlling modulator of the low-persistence phenotype prevalence, circumstantially supporting the conclusion that intercellular interactions during junctional trafficking are responsible for modulating PMN expression of GRK2.

## Discussion

PMNs are specialized immune cells circulating in the bloodstream that extravasate to migrate toward pathogens localized in the surrounding tissues. However, PMNs exhibiting fast, persistent migration often fail to contain bacteria growth *in vivo* (31). Instead, the ability to slow down and stop responding to chemoattractants seems to be essential for bacterial containment. Therefore, extravasated PMNs must exhibit plasticity in their migratory and phagocytic activity but the mechanisms that regulate this plasticity are still unclear. This study focused on how TEM, a crucial step of PMN extravasation, affects the migratory phenotype of PMNs and associated effector functions like phagocytosis.

Bayesian inference (26) of random walk in single cell trajectories yields time-varying values for two parameters representing each cell’s migratory persistence and random activity. Machine learning analysis of these two parameters for nearly 50,000 trajectories of PMNs migrating through 3D collagen matrices identified two distinct migratory phenotypes. One was characterized by fast, directed, chemotactically efficient *high-persistence* migration. The other was characterized by a slower, tortuous, *low-persistence* migration. Summary statistics like cell speed or chemotactic index have been used for decades to quantify the efficiency of PMN migration toward infection sites. Given the inherent variability of biological data, it is customary to average these statistics over multiple cells recorded under similar experimental conditions, assuming they all exhibit similar behavior. The heterogeneous migratory behavior of PMNs challenges this assumption and calls for caution when interpreting summary statistics computed over entire cell populations.

The low- and high-persistence migration phenotypes were present in dHL-60 cells across all different conditions studied and conserved in human blood PMNs. In line with this robustness, fast and slow subpopulations of neutrophils have been observed in the swarming locus of a murine ischemic-reperfusion injury model (43). More recently, we have reported heterogeneous migratory phenotypes in 3D-migrating T cells *in vitro* (44). Similarly, NK cells have been seen to intermittently exhibit in high- and low-persistence migratory modes depending on traction force exertion (20). The heterogeneity of migratory states is emerging as a preserved characteristic across different immune cells, further motivating the need to delineate how it is regulated since transitions between states could be a mechanism that endows PMNs with plasticity.

Our data suggest that TEM-associated stimuli promote a shift from the high- to the low-persistence migratory phenotype. TEM involves large PMN deformations and multiple molecular interactions with VECs that can cause mechanobiological responses (3, 8). For instance, dHL-60 cells traversing narrow constrictions experience chromatin reorganization (45), and prolonged cyclic deformations have been shown to depolarize PMNs (46). Experiments on human PMNs migrating up an fMLP gradient inside tapered microfluidic channels suggest that increasing mechanical confinement can trigger reverse chemotaxis (47). Reverse migratory behavior induced by physical constraints has been suggested to facilitate inflammation resolution (48).

In our experiments, crossing an empty membrane without VECs, an assay that only causes PMN deformation, predisposed dHL-60 cells to perform high-persistence migration contrary to our observations in VEC-lined membranes. This result suggests that heterotypic cell interactions during TEM could be significant modulators of PMN phenotype, overriding mechanical stimuli. Our experiments used 3-µm transwell membrane pores, as previously described (8), to allow enough cells to transmigrate and be collected for the 3D migration assay. Such pore size significantly exceeds the estimated endothelial junctional gap size (49), so our *in vitro* model could underrepresent PMN deformation. However, the transwell membrane material, polyethylene terephthalate (PET), is orders of magnitude stiffer than the VEC monolayer and the underlying basement membrane, which should compensate for the larger pore size.

We used a robust unsupervised clustering method to identify distinct migratory phenotypes (50, 51) based on parameterizing cell trajectories using a persistent random walk (PRW) model. The standard PRW model has been widely utilized for analyzing and simulating the migration of many cell types over flat surfaces (21–24) and has the convenience of relying on few parameters. However, applying this model to 3D cell migration has been more challenging (25). Studies relying on 4D live imaging produce hundreds of morpho-kinetic parameters per cell to help understand individual PMNs’ dynamic behavior under inflammation (43). However, phototoxicity, limited time resolution, and overall low throughput limit this approach to low numbers of cells that impede advanced statistical analyses. We utilized large-scale, label-free cell tracking to obtain multiple high-resolution trajectories per experimental video (4), incorporating a heterogeneous PRW model with time-dependent parameters that better captures the multi-scale patterns of 3D cell migration than the standard PRW model (26). An example of such patterns is the zigzagging described by cells belonging to the low-persistence cluster, which combines fast, highly-persistence segments with slower, quasi-random segments.

Although dHL-60 cells are a valid model system for many PMN studies (52), their imperfect differentiation efficiency and replication of PMN functions posit some limitations (41, 53). Therefore, we assayed PMNs obtained from human blood in our 3D migration device to validate our key findings in dHL-60 cells, e.g., the existence of two migratory phenotypes or the defining characteristics of each phenotype. Summary statistics like mean cell speed agreed with previously published results for dHL-60 cells and human PMNs (52). A more detailed comparison of cell trajectories based on the probability distribution of turning angles indicated a high level of similarity between human PMNs and the low-persistence cluster of dHL-60 cells. In particular, both cell types exhibited a clear secondary probability peak for large-angle turns, i.e., the hallmark of zigzagging. In this low-persistence cluster, dHL-60 cells from the ETW and CC conditions were most similar to human blood PMNs in terms of mean speed and CI, implying that TEM-mediated interactions with VECs could predispose dHL-60 cells to adopt a more physiologically representative phenotype. Also, PMNs from blood draws may have experienced diverse VEC interactions before collection, such as traversing the bone marrow endothelium or transmigrating from tissues back into the bloodstream (54, 55). These interactions could explain the similarity between human PMNs and ETW- and CC-conditioned dHL-60 cells.

GRK2 is an integrative signaling hub for modulating various aspects of cell migration processes (35). Upon chemokine stimuli, GRK2 constitutes serine/threonine protein kinases that specifically phosphorylate G protein-coupled receptors (GPCRs) to desensitize receptors (36). PMN swarming is a collective, self-amplifying response in which multiple PMNs form a perimeter around damaged tissue to prevent the spread of infection. GRK-2 deficient PMNs have exhibited reduced migration arrest and hence, ineffective swarm resolution (31, 56). Our experiments showed that GRK2 expression was enhanced in ETW-conditioned dHL-60 cells and that these cells had increased phagocytosis concurrent with a shift to the low-persistence phenotype. Moreover, GRK2 inhibition after ETW-conditioning blunted phagocytic activity and steered cells toward the high-persistence migratory phenotype. Therefore, our data suggest that TEM promotes a PMN phenotype with less persistent migration and increased phagocytic activity via upregulating GRK2 signaling.

It is possible that TEM stimulates a low-persistence migratory phenotype in PMN by degrading CXCR2 via GRK2 activation, consistent with evidence that GRK2 facilitates CXCR2 internalization and degradation (57). In agreement with this idea, we found that TW- and ETW-conditioned dHL-60 cells exhibited significant downregulation of CXCR2 surface expression. Moreover, our finding that CC conditioning did not change the expression level of CXCR2 suggests that the TEM-mediated modulation of this receptor could depend on mechanical stimuli. Low CXCR2 expression could be a natural modulator of PMN subpopulations. Indeed, a low-CXCR2-expression PMN cluster has been reported independently in different single-cell sequencing analyses (58, 59). Analysis of cell states showed that this cell cluster exhibits stronger anti-microbial activity than clusters with higher CXCR2 expression. Nevertheless, we cannot rule out that the molecular and surface changes of CXCR2 observed in our experiments result from other processes, including nuclear deformation-induced transcriptional changes or non-GRK2 activation-mediated CXCR2 expression. Experimental and analysis platforms such as the one presented here are poised to provide detailed, large-scale quantification of migratory behavior to help answer these questions.

Our study revealed that TEM-associated stimuli significantly affect the phenotype of PMNs. We showed increased phagocytic activity and concomitant loss of migratory persistence in PMNs after TEM, modulated by heterotypic interactions with VECs and mechanical deformation associated with junctional trafficking. Heterotypic interaction of dHL-60 with VECs recapitulates key migration features of primary PMNs in the low-persistence subpopulation, suggesting a more physiological relevant cell-line protocol of PMNs. Mechanical deformation during TEM is necessary for PMN functions in addition to heterotypic cell-cell interaction. GRK2 upregulation due to both effects from the TEM process is crucial for modulating PMN phenotype changes related to efficient pathogen identification and clearance. Immunosuppressants for treating chronic inflammation have numerous adverse side effects, including an increased risk of infections. Our findings on how the TEM process modulates PMN migration and functional plasticity may provide a rationale to identify specificity in targeting only one population of PMNs, thus offering opportunities for treating inflammation without compromising PMN’s anti-infection defense mechanisms.

## Materials and Methods

### Cell culture and differentiation

#### Neutrophil-like cell culture and differentiation

Cells from the human promyelocytic leukemia cell line (HL-60, ATCC) were cultured and differentiated into neutrophil-like cells (dHL-60) as previously described (60). Briefly, HL-60 cells were grown at 37C and 5% CO_2_ to an approximate density of 1×10^6^ cells/mL and passaged every two to three days by centrifugation and resuspension in Roswell Park Memorial Institute medium supplemented with l-glutamine (RPMI-1640, Gibco Thermo Fisher Scientific), 10% v/v fetal bovine serum (FBS, Omega Scientific), and 1% v/v penicillin-streptomycin (pen strep, Gibco Thermo Fisher Scientific). HL-60 cells were differentiated by taking approximately 1.5 x 10^6^ cells from a passage and resuspending them in supplemented RPMI-1640 with 1.3% v/v dimethyl sulfoxide (DMSO, Sigma) added. To ensure homogeneous distribution, the denser and more viscous DMSO was thoroughly mixed with the RPMI-1640 before adding cells. Experiments were conducted using dHL-60 cells cultured in the presence of DMSO between five and seven days. For experimental use in control (Ctrl) treatment conditions, day 5 dHL-60 cells were resuspended in supplemented M199 and incubated at 37C and 5% CO_2_ for thirty-six to forty-eight hours, then resuspended in fresh supplemented RPMI-1640 at the appropriate concentration for a given secondary assay.

#### Vascular endothelial cell culture

Primary human vascular umbilical vein endothelial cells (HUVECs, Cell Applications) were grown at 37C and 5% CO_2_ in Medium 199 with l-glutamine, 2.2 g/L sodium bicarbonate, and Earle’s salts (M199, Gibco Thermo Fisher Scientific) supplemented with 10% v/v FBS, 10% v/v Endothelial Cell Growth Medium (ECGM, Cell Applications), and 1% v/v pen strep until they formed a confluent monolayer. HUVEC were passaged following standard protocols for adherent mammalian cells: spent culture medium was aspirated, and cells were washed using Dulbecco’s phosphate buffered saline without calcium chloride or magnesium chloride (DPBS, Gibco Thermo Fisher Scientific); wash solution was aspirated and replaced with the pre-warmed 0.25% trypsin-EDTA (Gibco Thermo Fisher Scientific); the plate was incubated at 37C and 5% CO_2_ until at least 90% detachment was observed on an inverted microscope, at which point a volume of pre-warmed supplemented M199 equal to at least two times the volume of trypsin used was added to the plate and circulated by repeated pipetting to dislodge tenuously adherent cells; cells were collected via centrifugation, resuspended at the desired concentration in fresh supplemented M199, seeded in a situationally appropriate vessel, and incubated at 37C and 5% CO_2_ at least twenty-four hours prior to next use. Experiments were performed using cells between 4 and 7 passages (P4 – P7).

#### Primary neutrophil isolation from whole human blood

Primary neutrophils (PMNs) were isolated from whole human blood samples taken from healthy donors as previously described (61). Briefly, 7.5 mL of a red blood cell (RBC) sedimentation fluid comprising 3% v/v Gelatin, 0.9% v/v NaCl, and 0.1% v/v Dextrose was added to the bottom of a 50 mL conical tube at a volume appropriate to achieve a 1:2 ratio by volume with the 15 mL whole blood sample introduced on top. The conical was mixed by inversion three times, then incubated at room temperature for fifteen minutes until the RBCs sedimented and the solution stratified, leaving a layer of leukocyte-rich plasma at the top. That plasma was transferred into a 15 mL conical tube containing 3 mL of Ficoll-Paque Plus (Sigma), taking great care to avoid mixing the two solutions, and centrifuged at 4C and 1500 rpm for twenty minutes. After removing the supernatant, the pellet of RBCs and PMNs was resuspended in an RBC lysis solution comprising 1 mL DPBS and 5 mL H_2_O and mixed by inversion for forty-five seconds before adding 2 mL of 3% NaCl and centrifuging at 4C and 1500 rpm for five minutes. The supernatant was removed, and PMNs were gently resuspended in 1 mL of DPBS containing 1 g/L glucose for use in experiments.

#### Leukocyte TEM assay (ETW)

Transwell permeable supports were employed as the basis of an assay to replicate the impact of applied mechanical forces and intercellular interactions during transendothelial migration (TEM) on subsequent neutrophil physiology and migration mechanics (Figure 1A). Prior to loading, Transwell inserts with 3.0 μm pore polycarbonate membrane (12mm insert, Corning) were incubated with 200 μL of 50 ng/mL fibronectin (Millipore Sigma Aldrich) at 4C for at least twenty-four hours. The inserts were then washed twice with DPBS and equilibrated per manufacturer recommendation by incubating them at 37C and 5% CO_2_ for twenty-four to forty-eight hours with 1 mL of fresh supplemented M199 in the lower chamber and 500 μL in the upper chamber.

Following equilibration, the medium in the lower chamber was replaced with 1 mL fresh supplemented M199, and the medium in the upper chamber was replaced with 500 μL HUVEC suspended at a concentration of 4.5 x 10^5^ cells/mL in supplemented M199. Inserts were incubated at 37C and 5% CO_2_ for thirty minutes, then washed with supplemented M199 to remove HUVEC that were not adhered to the polycarbonate membrane and incubated for twenty-four hours to form a confluent monolayer. The medium was then aspirated from the upper chamber and replaced with fresh M199 with 20 ng/mL tumor necrosis factor alpha (TNF-α) for at least eight hours to activate the HUVEC monolayer.

Following TNF-α activation, the medium in the lower chamber was refreshed, day 5 dHL-60 cells were resuspended in supplemented M199 and loaded into the upper chamber at a density of approximately 8 x 10^6^ cells, and the inserts were incubated at 37C and 5% CO_2_. After thirty-six to forty-eight hours, dHL-60 cells that had migrated through the HUVEC monolayer and the polycarbonate membrane, termed ETW treated cells, were collected from the lower chamber and resuspended in fresh, supplemented RPMI-1640 at an appropriate concentration for further pre-treatment or use in experiments.

#### Empty transwell-based leukocyte treatment assay (TW)

To decouple the impact of physical interaction with and deformation required to pass through the membrane in the TW support from biochemical and physical interactions with the HUVEC monolayer on subsequent leukocyte interstitial migration, an empty transwell (TW) treatment condition was employed (Figure 1C). Untreated inserts were equilibrated with supplemented M199 as described above, then loaded with day 5 dHL-60 cells and incubated at 37C and 5% CO_2_. After thirty-six to forty-eight hours, cells that had migrated through the polycarbonate membrane of the TW support were collected from the lower chamber of the insert and resuspended in fresh supplemented RPMI-1640 at an appropriate concentration for further treatment or use in experiments.

#### HUVEC and dHL-60 co-culture assay (CC)

To further decouple the influence of intercellular biochemical signaling and intimate physical contact between HUVEC and dHL-60 cells during TEM from the large-scale cell body deformations and external forces applied to dHL-60 cells as they traverse micron-scale junctional gaps in the HUVEC monolayer, we introduced a co-culture (CC) treatment condition. HUVEC were cultured to confluence in a 10 cm plate, then activated with TNF-α as previously described. Then, the spent medium was aspirated and replaced by day 5 dHL-60 cells suspended in supplemented M199 at approximately 1.6 x 10^7^ cells/mL, and the plate was incubated at 37C and 5% CO2. After thirty-six to forty-eight hours, the medium, containing cells in suspension, was circulated by repeated pipetting to dislodge any dHL-60 cells still loosely adhered to the HUVEC monolayer, then collected and resuspended in supplemented RPMI-1640 at a concentration appropriate for use in further experiments.

#### Multi-color flow cytometry

Samples were prepared for a four-color flow cytometry (FCS) panel following standard protocols for antibody staining. For each experimental condition, four technical replicates were prepared for the full FCS panel in addition to four technical replicates, each of fluorescence minus one control (all fluorophores in the panel except one, to assist with gating), and single stain controls (to assist with compensation). Unstained controls were resuspended in PBS and run without any additional treatment. Unless otherwise specified, antibodies were purchased from Abcam. For ICAM-1 and CD11b staining, cells were collected, resuspended in 10 μL of DPBS, mixed with the appropriate primary antibody (goat ICAM-1 or rabbit CD11b) at a 1:1 (v/v) ratio, and incubated on ice for 30 – 60 min. CXCR1 and CXCR2 stains were purchased as pre-conjugated antibodies and did not require this step. Samples were then resuspended in 10 μL of fresh DPBS, mixed with the appropriate fluorophore-conjugated secondary antibody or pre-conjugated stain at a 1:1 (v/v) ratio, and incubated covered on ice for 45 – 60 min. Samples were then washed once with DPBS and resuspended in 300 μL of DPBS for experimental use. 30,000 events were collected for each sample using a BD Accuri C6 Plus personal flow cytometer. Standard compensation, gating, and subsequent analysis were performed using FlowJo.

#### Total RNA extraction using TRIZOL reagent

RNA isolation was performed following the TRIZOL RNA isolation protocol from the W.M. Keck Foundation Biotechnology Microarray Resource Laboratory at Yale University. Briefly, the cells of interest were resuspended in ice-cold DPBS, lysed via repetitive pipetting with TRIZOL reagent (Ambion; 1mL reagent per 5-10 x 10^6^ cells), and allowed to stand at room temperature for 5 min to ensure complete disassociation. The suspension was centrifuged to remove any cell debris, and then the supernatant was transferred to a new tube. Chloroform was added to the supernatant at a ratio of 0.2 mL for every 1 mL of TRIZOL used, vortexed vigorously, and left to stand at room temperature for two min before centrifuging to generate complete phase separation. The clear, upper aqueous phase was transferred to a new tube.

RNA was precipitated from the solution by adding isopropyl alcohol to the aqueous phase at a ratio of 0.5 mL for every 1 mL of TRIZOL used, incubated for 10 min at room temperature, and centrifuged to pellet the RNA precipitate. After removing the supernatant, the RNA precipitate was washed twice with a volume of 75% v/v ethanol greater than or equal to the volume of TRIZOL used, air-dried for five to ten minutes, and redissolved into 100 μL of DEPC-treated water. RNA concentration was measured using a Nanodrop 2000c (Thermo Scientific).

#### First-strand cDNA synthesis

First-strand cDNA was synthesized using M-MLV reverse transcriptase (RT; Invitrogen) following the protocol included with the product information sheet. Briefly, for a standard 20 μL reaction volume, 1 μL of oligo DT primers, a volume of resuspension containing 1 ng to 5 μg of total RNA isolated as described in the previous section, 1 μL of 10 mM dNTP mix were added to a nuclease-free microcentrifuge tube with sterile water such that the total volume was 12 μL. The tube was heated to 65C for five min in a MyCycler thermal cycler (BioRad) and chilled on ice before adding 4 μL of 5x first-strand buffer, 2 μL of 0.1 M DTT, and 1 μL of RNaseOUT recombinant ribonuclease inhibitor (40 units/μL). Contents were mixed and incubated in the thermal cycler at 37C for two min before adding 1 μL of M-MLV RT and returning the tube to the thermal cycler, where the mixture was held at 37 C for fifty min, immediately followed by fifteen min at 70 C. cDNA was either used immediately as described below or stored at −80C until needed.

#### Real time quantitative polymerase chain reaction (RT-qPCR)

The newly synthesized first-strand cDNA was used as a template for amplification in RT-qPCR following the protocol included in the product information sheet for iQ supermix (BioRad). Briefly, assay master mix was prepared by mixing 7.5 μL of PowerTrack SYBR green master mix (2x, Applied Biosystems) with 100 nM each of the forward and reverse primers (IDT, see Table 1 for sequences) and a volume of sterile water such that, when the master mix was added to a volume of RT product containing 100 ng of cDNA, each well of the qPCR plate contained a total reaction volume of 15 μL. The total volume of master mix prepared was appropriately scaled to account for technical triplicates of each gene of interest for each biological repeat and treatment condition.

**Table 1:**
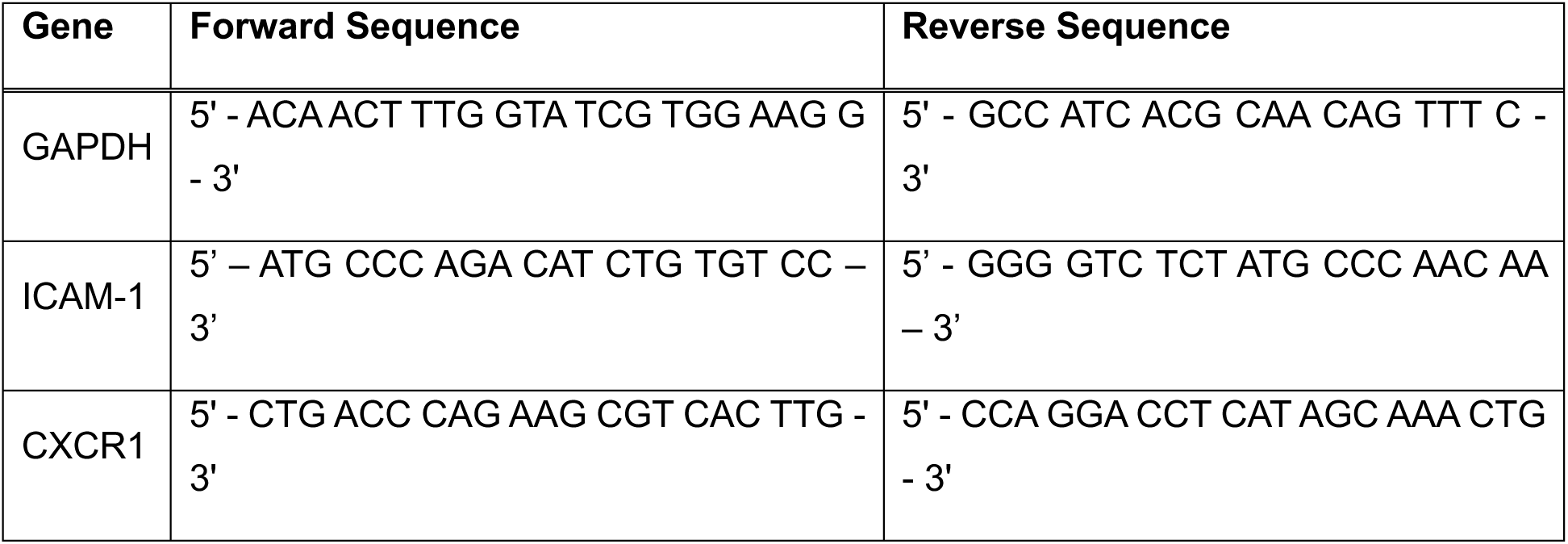
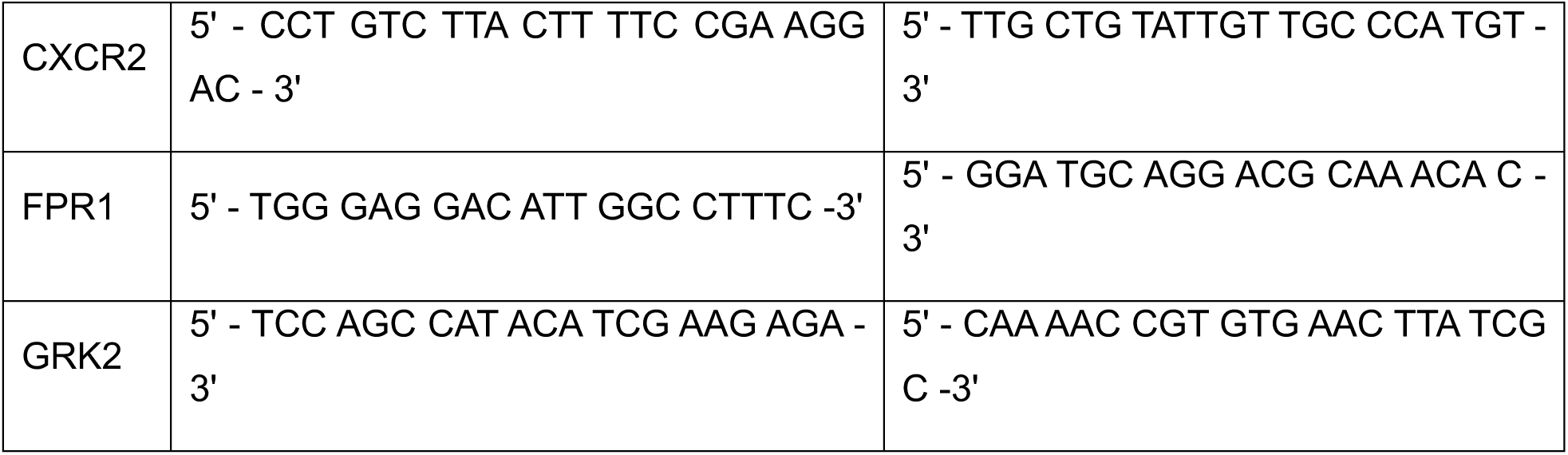
Forward and reverse sequences of oligos used for RT-qPCR.

Once each well was loaded, the plate was sealed with an optically transparent film, vortexed for 30 seconds to ensure thorough mixing of the reaction components, loaded into a CFX Connect Real-Time System (BioRad), and run following the manufacturer’s thermal cycling conditions. Specifically, the reaction plate was held at 50 C for 2 minutes to activate uracil DNA glycosylase, then at 95 C for 2 minutes to activate DNA polymerase, before repeating the denaturing, annealing, and extending steps (95 C for 15 seconds, 58.1 C for 15 seconds, and 72 C for 1 minute, respectively) for 40 cycles.

The resulting data was analyzed in Microsoft Excel using the comparative C_*t*_ method (62). Briefly, expression levels for a gene of interest were normalized to a comparative baseline within each biological replicate by taking the difference between the C_*t*_ value for that gene and the C_*t*_ value of the housekeeping gene GAPDH (ΔC_*t*_). ΔC_*t*_ values are expressed in relation to the control case for the purpose of comparison across treatment conditions by taking the difference between ΔC_*t*_ for a condition of interest and ΔC_*t*_ for the control case (ΔΔC_*t*_). Figures were generated using GraphPad Prism, showing log fold change compared to control, or 2^−ΔΔC_T_^, for ease of visualization. Statistically significant changes were determined using Kruskal-Wallis test for multiple comparisons, performed on ΔC_*t*_ values using GraphPad Prism. Between-run variation resulting from the use of multiple qPCR plates to amplify the same cDNA was removed by applying a previously reported mathematical transformation as part of the Excel analysis pipeline (63).

#### *In vitro* 3D directed migration assay

*In vitro* assays of 3D directional migration were performed in a custom-built chemotaxis device (Figure 2A) as previously described (4). Briefly, 25 mm x 25 mm No. 1.5 coverslips (Corning) were placed on a 110 C hot plate and treated on one side with 300 μL of 0.1 M NaOH with 0.02% v/v Triton X. Once the NaOH solution evaporated, the coverslips were washed in distilled H_2_O and allowed to dry. Then 50 μL of (3-aminopropyl) triethoxysilane (Sigma-Aldrich) was added to the treated surface of the coverslip for 5 min. Coverslips were washed as described above, and the treated surface was exposed to 100 μL of 0.5% v/v glutaraldehyde for 30 to 120 min. Coverslips were allowed to air dry overnight in a vented, covered container or in a 65 C oven for 1 hour in the same vented, covered container. Next, a 12-mm-diameter hole was punched in the center of a 50 x 9-mm Falcon petri dish (BD Falcon) and attached a treated glass coverslip, treated side facing up, to the bottom of the petri dish to create a glass bottom dish. A second treated coverslip was then attached to the top side of the dish, treated side facing down and occluding approximately three quarters of the hole to create a pocket of space in the center of the hole that has silane-treated glass on both sides and a small loading port at the top. Both coverslips were attached with vacuum grease (Beckman Vacuum Grease Silicone), and all edges in contact with the petri dish were further sealed with clear quick dry nail polish (Sally Hansen Insta Dri). Before gel fabrication, devices were equilibrated at 37C and 5% CO_2_ for a minimum of thirty minutes to help mitigate bubble formation during loading.

#### Collagen gel fabrication and device loading

Collagen gels were fabricated by neutralizing a solution of rat-tail type-I collagen monomers dissolved in a 0.2 N acetic acid (Corning) with 1 N NaOH in proportions recommended by the manufacturer’s alternate gelation protocol to generate gels with a final collagen concentration of 1 mg/mL. A suspension of supplemented RPMI-1640 containing approximately 7.5 x 10^5^ dHL-60 cells was used in place of the recommended water. All constituents for gel fabrication were kept, mixed, and degassed on ice to prevent premature self-assembly, and the mixture was degassed for 5 min prior to loading. Once prepared, 113 μL of the collagen gel solution was pipetted into the pocket of each device. Devices were then incubated vertically at 37C and 5% CO_2_ for thirty minutes to allow the gel to self-assemble in the lower half of the pocket. The remaining volume of the pocket was filled with 113 μL of supplemented RPMI-1640 with 50 nM fMLP and immediately imaged.

#### Image acquisition and cell tracking with bulk drift correction

Bright-field images for population migration experiments were obtained using an enclosed Leica DM16000 B microscope with a 5x air lens at 37C and 5% CO_2_, with the built-in magnifier retracted to produce images more amenable to automated cell tracking. Four planes, evenly spaced 200 μm apart to ensure the same cells were not captured in multiple planes, were imaged within each collagen gel every Δt = 10 sec.

Automated cell tracking for standard time resolution experiments was performed using the Fiji plugin Trackmate (64, 65), discarding the first 50 frames of the time course for each experiment to eliminate migration artifacts caused by rapid gel drift. Cells were identified using a Laplacian of Gaussian (LoG) edge detection filter with median filtering enabled, and frame linking was performed using a linear assignment problem (LAP) tracker. Final cell tracks were filtered based on quality and duration, then exported to MATLAB for bulk drift correction and further analysis.

Drift velocity was calculated based on the bulk motion of non-motile cells within each experiment. Briefly, the first and second moments of inertia were calculated for each cell track and used as the basis for selecting the 15% of the experimental population with the lowest overall motility. Visual inspection confirmed that these cells were non-motile, and the one-minute smoothed median velocities of the non-motile population in each Cartesian direction were taken as the bulk drift velocity for the experiment. The drift velocity was then subtracted from each cell track within the experimental population to correct for bulk gel deformations during the experiment.

#### Summary migration statistics with respect to time

Cell centroids obtained from the tracking algorithm described above were used to calculate average instantaneous cell speed over Δt = 10 sec imaging intervals,

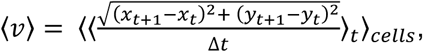

then converted to units of microns per minute. Average chemotactic index (CI) values were computed by taking the cosine of the angle *θ* between the chemotactic gradient vector and the displacement vector of each cell, *CI* = 〈cos *θ*〉_*cells*_.

#### Analysis of trajectories in intrinsic coordinates

The shape of cell trajectories was analyzed in terms of distance traveled and angular orientation (intrinsic coordinates) as previously described (4), rather than in terms of time to eliminate the confounding influence of variations in migration speed between conditions. The distance travelled by a cell versus time was first computed as 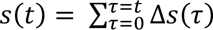 where 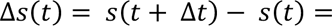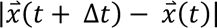. This information was used to obtain 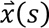 via interpolation (interp1, MATLAB). The vector tangent to the trajectory was obtained as 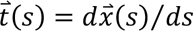. The mean squared displacements (MSDs) were computed as a function of distance lags, Δ*s*, i.e., 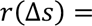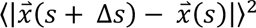, where the bracket denotes an ensemble average for all the cells in each experiment and along all values of *s* along each trajectory. Similarly, we computed the autocorrelation function for the vector tangent to the trajectory, 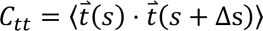. Changes in orientation of 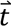 with respect to Δ*s* were calculated as, 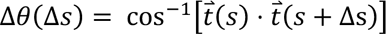, in addition to their PDF conditioned to Δ*s*, *p*(*θ*| Δ*s*).

#### Estimation of migratory persistence length

Based on the presence of two distinct decay regimes evident in semi-log plots of *C_tt_*, the migratory persistence length for each condition was estimated by fitting *C_tt_* to a double exponential decay model, expressed as 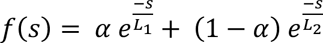 for each treatment condition (lsqcurvefit, MATLAB). The double exponential fit requires that each condition be described in terms of two characteristic lengths, *L*_1_ and *L*_2_, defined so that *L*_1_ > *L*_2_. In all experiments, *L*_2_ characterized a rapid autocorrelation decay, with average characteristic length <2 mm. Thus, the migratory persistence length was taken to be *L_P_* = *L*_1_. Error bars were calculated via bootstrapping. Briefly, 25% of the cells in a condition were randomly selected for removal from the population and *L_P_* was calculated for that subpopulation of cells. This process was repeated twenty times. Mean and SEM were then calculated based on the twenty *L_P_* values.

#### Phagocytosis assay

To measure changes in phagocytic behavior of dHL-60 cells before and after TEM, dHL-60 cells were cultured and differentiated as described in (cell culture section), and ETW sorted cells were prepared by performing the treatment assay described in (cell culture section). dHL-60 cells in the condition of interest were resuspended with pHrodo Green Zymosan Bioparticles conjugate for phagocytosis (Thermo Fisher Scientific) at the manufacturer’s recommended concentration and co-incubated at 37C and 5% CO_2_ for 30 min. The sample was then resuspended in 300 μL of DPBS. pHrodo particles are pH sensitive and will only emit fluorescence in highly acidic intercellular environments such as endosomes and lysosomes. As such, we could quantify the number of cells that had performed phagocytosis by employing single-color flow cytometry following the measurement and analysis protocols outlined in (flow cytometry section).

#### Inhibition of GRK2 expression in dHL-60 cells

GRK2 inhibition experiments were performed using CMPD101 (Tocris Bioscience). For control conditions, day 5 dHL-60 cells were resuspended in supplemented M199 and incubated at 37C and 5% CO_2_ for thirty-six to forty-eight hours. They were then resuspended in supplemented RPMI-1640 with 30 μM CMPD101, incubated for 30 min at 37C and 5% CO_2_, then resuspended in fresh supplemented RPMI-1640 at an appropriate concentration for experimental use. For TW sorted conditions, cells were collected from the lower chamber of the sorting assay, as described above, resuspended in supplemented RPMI-1640 with 30 μM CMPD101, incubated for 30 min at 37C and 5% CO_2_, then resuspended in fresh supplemented RPMI-1640 at an appropriate concentration for experimental use.

#### Time-dependent Bayesian sequential inference of heterogenous random walk

A 2D, first-order, auto-regressive process was used to model the persistent random walk of individual cells with time-varying motility parameters. The time series of the velocity vector of an individual cell, {***v***_1_, …, ***v***_*t*−1_, ***v***_*t*_, ***v***_*t*+1_, …, ***v***_*N_t_*−1_} was numerically obtained with 2^nd^ order finite differences, where *N*_*t*_ is the number of images in the timelapse. Adapting the methodology of Metzner, Mark (26), the stochastic change in the velocity of a cell given the previous time step is described by the time-discrete stochastic process,

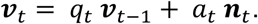

Here, *q_*t*_* ∈ [−1,1] corresponds to the local persistence of the random walk, *a_*t*_* ∈ [0, ∞) represents the local activity (stochastic amplitude) that sets the spatial scale of the random walk and ***n****_*t*_* represents an uncorrelated, normally distributed noise with unit variance. The persistence *q_*t*_* is a measure of the directional correlation between the successive time steps, not necessarily in the direction of the chemotactic gradient. *q_*t*_* = 1 corresponds to a straight line and *q_*t*_* = 0 to a non-persistence Brownian diffusion. Higher values of *a_*t*_* correlate with increase in metrics such as cell speed, total displacement and track length. Together, the activity and persistence parameters determine the variance of displacement steps within a cell track according to *var*(*v*) = *a*^2^/(1 – *q*^2^) independently in each coordinate. Bayesian sequential inference (adapted from Metzner, Mark (26)) was performed in MATLAB to estimate the time-varying heterogeneous migration parameters of each individual cell.

#### Unsupervised identification of heterogenous sub-population from the random walk characteristics

Each individual cell was analyzed for changes in the migratory state during the duration of imaging to avoid averaging over different motility modes. A density-based spatial clustering (DBSCAN) algorithm (50) in MATLAB was employed to identify outliers and clusters in the inferred random walk parameters (*q_*t*_*, *a_*t*_*) within a single cell trajectory data. The parameter *a_*t*_* for each cell was normalized to the range [0,1] for subsequent analyses. The parameters *ε* = 0.1 and *minPts* = 10 was used for the DBSCAN algorithm. This procedure is termed ‘local clustering’. If more than one cluster was detected, they were treated as individual cells for further analysis. Each migrating cell’s trajectory was characterized by its average random walk parameters 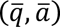 of persistence and activity after local clustering.

To automatically identify sub-populations and shifts in the sub-populations across all experimental conditions (e.g. control, pharmacological treatments, etc.), cell-averaged random walk parameters 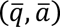 were pooled together. A Variational Bayesian Gaussian Mixture Model (VB-GMM) was used to cluster in the parameter space of 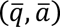. A Gaussian mixture model has the advantage of identifying elliptical clusters with anisotropic variances and the Bayesian formulation allows for automatically inferring the number of clusters from the data. This step, termed ‘global clustering’, was performed to autonomously identify heterogenous sub-populations based on the trajectory data alone.

The VB-GMM clustering algorithm fitted the datapoints in cell-averaged random motility parameter space with *K* Gaussian distributions delineating the densely populated regions (51), using code adapted from mixGaussVb.m in the MATLAB File Exchange, noting that we corrected an error in the posterior covariance matrix in the code. The GMM model assumes a mixture of *K* Gaussian clusters, each with a mean *μ_k_* and precision matrix Λ*_k_*, such that the distribution of the cell-averaged random walk parameters 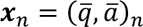 is

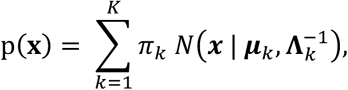

where π*_k_* is the mixing coefficient, such that 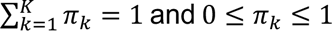, controlling the proportion of each cluster. A symmetric Dirichlet distribution was chosen as the conjugate prior for π*_k_* with the density,

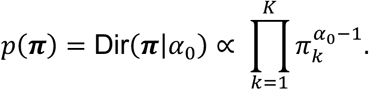

The concentration hyper-prior *α*_0_ controls the sparsity of cluster numbers and the influence of data on the inference of the number of components *K*. *α*_0_ values closer to 0 impart *a priori* belief that prefers fewer clusters that is influenced by data and as *α*_0_ → ∞, the posterior of *π_k_* → 1/*K* (no sparsity). For this manuscript, *α*_0_ = 1 was used which produces a flat Dirichlet distribution letting the parameter *K* be determined only from the data. Such a flat distribution is advantageous when working with many data points. For the parameters *μ_k_* and *Λ_k_*, a conjugate Gaussian-Wishart prior with zero mean was prescribed. The variational inference of the GMM mean *μ_k_* and precision matrix *Λ_k_* as well as the number of clusters *K* was based on factorized approximation to the posterior distribution through a latent variable representation. The specific details of the inference method are omitted here, and the reader is referred to Bishop *et al* (51).

